# Laminar dynamics of beta bursts in human motor cortex

**DOI:** 10.1101/2021.02.16.431412

**Authors:** James J Bonaiuto, Simon Little, Samuel A Neymotin, Stephanie R Jones, Gareth R Barnes, Sven Bestmann

## Abstract

Modulation of motor cortical activity in the beta frequency range is one of the strongest and most studied movement-related neural signals. At the single trial level, beta band activity is often characterized by transient bursting events rather than slowly modulating oscillations, suggesting a more rapid, information-encoding functional role than previously believed. Insight into how beta bursts are generated in sensorimotor circuits can provide important constraints to theories about their functional role for movement control. To this end, we leverage and extend recent developments in high precision MEG for temporally resolved laminar analysis of burst activity, combined with a neocortical circuit model that simulates the biophysical generators of the electrical currents which drive beta bursts. This approach pinpoints the generation of beta bursts in human motor cortex to distinct excitatory synaptic inputs to deep and superficial cortical layers, which drive current flow in opposite directions. These laminar dynamics of beta bursts in motor cortex align with prior invasive animal recordings within the somatosensory cortex, and suggest a conserved mechanism for somatosensory and motor cortical beta bursts. More generally, we demonstrate the ability for uncovering the laminar dynamics of event-related neural signals in human non-invasive recordings.

## Introduction

One of the most conspicuous movement-related brain signals is motor cortical activity in the beta frequency band (13-30Hz). It is now well established that trial-averaged sensorimotor beta activity decreases prior to movement and increases following movement (Cassim et al., 2001; Jasper and Penfield, 1949; Jurkiewicz et al., 2006; Kilavik et al., 2013; Pfurtscheller, 1981; Pfurtscheller et al., 1996; Pfurtscheller and Lopes da Silva, 1999). This gradually modulating average signal has been linked to a wide range of functions (Cao and Hu, 2016; Cassim et al., 2001; Chandrasekaran et al., 2019; Engel and Fries, 2010; Jasper and Penfield, 1949; Khanna and Carmena, 2017; Kilavik et al., 2013; Pfurtscheller, 1981; Pfurtscheller et al., 1996; Sanes and Donoghue, 1993; Spitzer and Haegens, 2017; Tan et al., 2016, 2014), but its functional role remains undetermined.

Moreover, at a single trial level, beta activity does not appear to be a sustained oscillation but instead is dominated by transient bursting events (Feingold et al., 2015; Jones et al., 2009; Little et al., 2019; Murthy and Fetz, 1996, 1992; Sherman et al., 2016; Shin et al., 2017; Spitzer and Haegens, 2017). The probability of burst occurrence decreases before, and peaks after the movement, thus giving the appearance of slow beta amplitude modulation after averaging these bursts over multiple trials (Jones, 2016; Little et al., 2019; Seedat et al., 2020).

It is becoming increasingly clear that beta bursts recorded over sensorimotor cortices have an information-encoding role in movement control. For example, the rate and timing of beta bursts in humans are more predictive of behavior than mean beta amplitude (Hannah et al., 2020; Jana et al., 2020; Little et al., 2019; Shin et al., 2017; Wessel, 2020). Moreover, symptom severity in Parkinson’s disease is correlated with burst amplitude and duration (Cagnan et al., 2019; Tinkhauser et al., 2018, 2017b; Torrecillos et al., 2018), linking burst abnormalities to pathological movement control and motivating burst activity as a target for clinical applications of deep brain stimulation (Little et al., 2013; Moraud et al., 2018; Tinkhauser et al., 2017a). While the rich spatial and temporal structure unveiled by the discovery of beta bursts provides an opportunity to more precisely identify their relevance for healthy and pathological movement, this is tempered by a lack of understanding of the mechanisms underlying the generation of sensorimotor beta bursts.

Theories of cortical computation have proposed distinct roles for neural activity in various frequency channels and projections originating from deep and superficial cortical layers (Adams et al., 2013; Arnal and Giraud, 2012; Bastos et al., 2018, 2015, 2012; Donner and Siegel, 2011; Fries, 2015, 2005; Friston and Kiebel, 2009; Jensen et al., 2015; Jensen and Mazaheri, 2010; Stephan et al., 2017; Wang, 2010), but relatively few of these theories address the generation of frequency-specific activity by inter-laminar dynamics within cortical circuits. Recent computational neural models of beta burst generation (Law et al., 2019; Neymotin et al., 2020; Sherman et al., 2016) suggest that such bursts can be generated by strong excitatory synaptic input to superficial cortical layers lasting ~100ms, temporally aligned with a broad, weaker input to deep layers lasting ~50ms. These synchronized inputs cause current to propagate in opposite directions within a cortical column, resulting in a cumulative dipole with the stereotypical wavelet shape in the time domain as measured by local field potentials (LFPs), electroencephalography (EEG), and magnetoencephalography (Karvat et al., 2020; Kosciessa et al., 2020; Little et al., 2019; Sherman et al., 2016). The spectral frequency of the burst is determined by the duration of the strong supragranular synaptic input that drives current towards deep layers of the cortex, generating the prominent ~50ms peak in the beta burst wavelet shape. The surrounding tails of the beta burst peak emerge from synaptic input to the deep layers that drive current flow toward superficial layers. Laminar current source density recordings from primary somatosensory cortex in rodents and non-human primates provided initial support for these model-derived predictions (Sherman et al., 2016).

Until recently, testing such predictions noninvasively in the human brain was constrained by the low temporal resolution of fMRI and the low spatial precision of EEG and MEG. However, recent developments in high precision MEG (hpMEG; Little et al., 2018; Meyer et al., 2017; Troebinger et al., 2014b) have demonstrated sensitivity of the recorded signals to the orientation of cortical columns (Bonaiuto et al., 2020), and the ability to test hypotheses concerning the laminar dominance of frequency-specific induced activity (Bonaiuto et al., 2018a, 2018b). We here extended these techniques to develop a temporally resolved laminar analysis which yields estimates of the relative strength of superficial and deep cortical layer activity over the time course of pre- and post-movement beta bursts.

To validate the ability to recover *a priori* known patterns of laminar activity, we first used a detailed biophysical model of beta burst generation to create simulated MEG data using a generative MEG model. We then tested the ability of the new laminar analysis to recover the simulated strong superficial layer activity at the peak of the burst and deep layer activity surrounding the peak. We then generated predictions for sensor level activity and laminar dynamics from a range of alternative synthetic models with varying dipole laminar locations and current flow directions.

When applying these analyses to human MEG data, we found that the laminar profile of both pre- and post-movement beta bursts in motor cortex conformed to the biophysical model’s predictions: activity surrounding the burst peak localized to deep cortical layers, whereas the peak corresponded to activity predominantly in superficial layers. When compared against alternate models of burst generation, we found support only for the model in which bursts are generated by distinct synchronous inputs to deep and superficial cortical layers, which in turn drive current flow in opposite directions. These results thus validate the predictions of the biophysical model in human motor cortex, support a rapid and dynamic functional role for sensorimotor beta bursts, and demonstrate the possibility for uncovering the laminar specificity of dynamic neural signals in non-invasive human MEG recordings.

## Results

### Beta bursts in human motor cortex have a stereotyped waveform shape

We first identified bursts of precentral beta activity in MEG data from human participants performing a visually cued action decision-making task involving button press responses made using the right hand (Bonaiuto et al., 2018a). Bursts were identified at the source level as periods when beta amplitude in the hand area of left motor cortex exceeded an empirically defined threshold (Little et al., 2019). We segmented each participant’s dataset into 200ms epochs, centered on the peak of each burst, and split each dataset into bursts that occurred prior to (M=326.8 SE=133.3 bursts per subject), or after (M=722.5, SD=155.0 bursts per subject) the button press response in each trial. After alignment, the average time series of pre-movement beta bursts matched the stereotyped wavelet-like shape previously described in somatosensory cortex (Sherman et al., 2016), and appeared as a single dipolar field pattern centered over left sensorimotor cortex with a transient reversal in direction around the burst peak (**Figure 1**A). This mean pre-movement burst waveform shape was consistent across all participants (**Figure 1**B). Beta bursts that occurred post-movement had the same spatial and temporal features as pre-movement bursts (**Figure 1C**), but had a greater peak amplitude (*X^2^*(1)=95.77, *p*<0.001; **Figure 1**D). This suggests that a common mechanism may underlie the generation of pre- and post-movement motor beta bursts.

**Figure 1.**
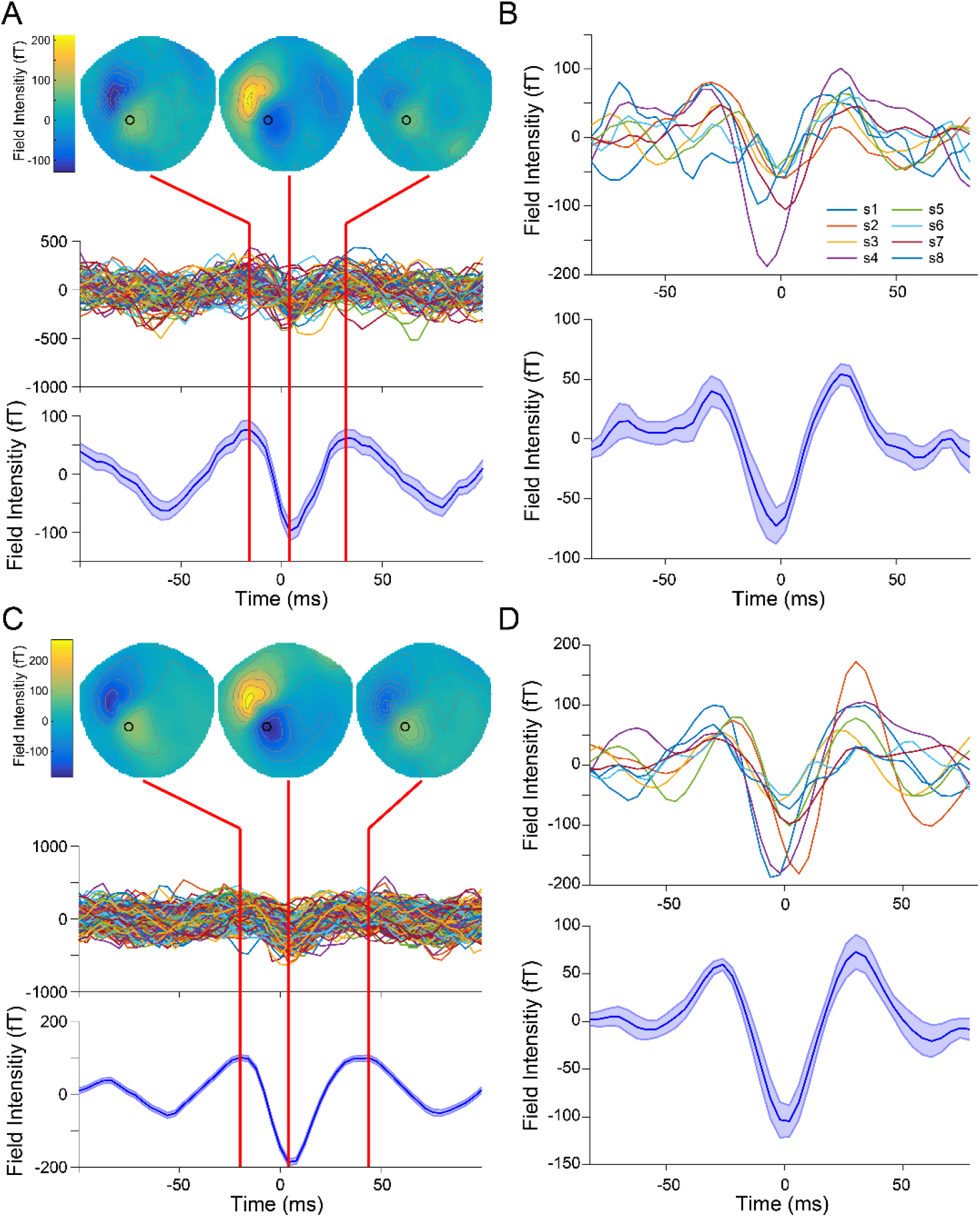
Sensor-level pre- and post-movement beta bursts. A) Single subject sensor-level data aligned to the peak of pre-movement motor beta bursts. The mean spatial topography (top) indicates a field reversal over left sensorimotor cortex centered on the burst peak. Individual premovement burst waveforms (middle), taken from the sensor indicated by a black circle in the spatial topography (MLP34), exhibit a stereotyped, wavelet-like shape that becomes more apparent when averaged over all premovement bursts (shaded area indicates the standard error of the burst waveform across all bursts). B) Mean pre-movement burst waveforms from all subjects after alignment with a Woody filter (top) and averaged over subjects (bottom). The shaded area indicates the standard error of the mean burst waveform across subjects. Post-movement motor beta bursts have the same spatial and temporal pattern as pre-movement bursts at the single subject (C) and group level (D).

### Biophysical computational modeling predicts a specific laminar mechanism for beta burst generation

We next sought to determine the putative layer-resolved cortical mechanisms behind the generation of beta bursts in motor cortex. For this purpose, we extended previously developed hpMEG techniques (Bonaiuto et al., 2018a, 2018b) to yield a temporally resolved estimate of laminar activity. This new analysis involved construction of generative MEG models of oriented current patches from white matter and pial surfaces extracted from each subject’s MRI volume (**Figure 2**A), representing deep and superficial cortical layers. Each subject’s burst-aligned data were then averaged, and source inversion was conducted on the average beta burst time series to localize the bursts in source space (**Figure 2**B). After determining the source space locations of precentral beta bursts, we then proceeded to use a sliding time window analysis to compare the relative strength of current flow in deep and superficial layers over the time course of the bursts. At each pial surface vertex selected by the localizer inversion, we identified the corresponding white matter surface vertex in the direction of the estimated orientation of the cortical column at that location using the link vectors method (Bonaiuto et al., 2020). These vertex pairs were then used as Bayesian priors in a sliding time window source inversion (**Figure 2**C). This involved determining the more likely generative (pial and white matter) model, given the burst activity, within a small time window of the average burst waveform, using Bayesian model evidence (approximated by free energy). Following each time window analysis, we advanced the time window to obtain a time series of the free energy difference between the models (approximating the Bayes factor; **Figure 2**C). The resulting time series provides an estimate of the likelihood that the electrical currents generating the field signal were stronger either in deep or superficial layers (as approximated by our pial and white matter surfaces) at each time point.

**Figure 2.**
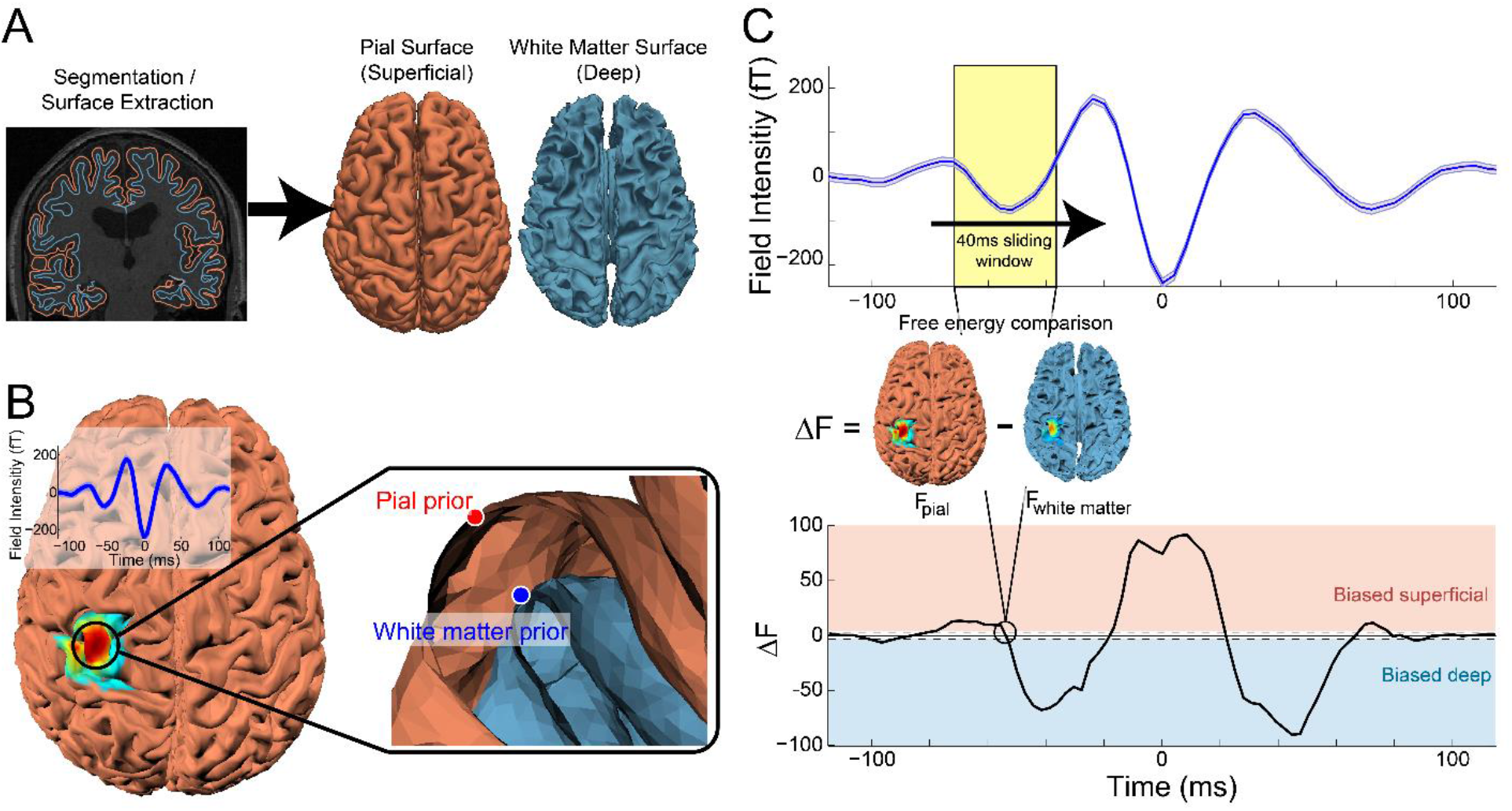
Sliding time window laminar source inversion. A) Pial and white matter surfaces are extracted from quantitative maps of proton density and T1 times from a multiparameter mapping MRI protocol. B) Source inversion over the entire burst time course was used to localize the average beta burst sensor signal (inlay). The peak pial surface vertex and the corresponding vertex on the white matter surface were used as priors in the following sliding window inversion. C) Sliding time window source inversion was performed using a 40ms wide window. For each iteration, source inversion was run using a pial generative model with the pial vertex from the localization inversion as a prior, and using a white matter generative model with the corresponding white matter vertex as a prior. The difference in free energy between the two models (F_pial_ – F_white matter_) was used to determine the laminar locus of dominant activity as the window advanced along the average time series of the beta burst (example data shown from a single human subject).

Before applying this analysis to human MEG data, we first verified that this approach could reliably identify the time course of laminar activity in simulated data for which the ground truth was known. We therefore simulated a set of beta bursts using the biophysical model, as in our previous work (**Figure 3**A, Sherman et al., 2016). In this model, the synaptic drive is generated by trains of action potentials in predefined temporal patterns, whose Gaussian shaped histograms are schematically depicted in **Figure 3**A, that activate fast excitatory synapses at layer specific locations as shown. This drive generates intracellular current flow up or down within the cortical pyramidal neuron dendrites, and the cumulative current dipole moment is estimated from the net longitudinal intracellular currents across the pyramidal neurons, multiplied by their length (Neymotin et al., 2020; Sherman et al., 2016; see Methods for further details). When only the superficial distal drive is present, the cumulative current dipole exhibits a downward deflecting current into the cortex (**Figure 3**B, top), and when only the deep proximal drive is present, the cumulative current dipole exhibits an upward deflecting current out of the cortex (**Figure 3**B, bottom). The cumulative dipole moment from the combination of these inputs results in a dipole moment that resembles the recorded beta burst waveform (**Figure 3**B, middle).

**Figure 3.**
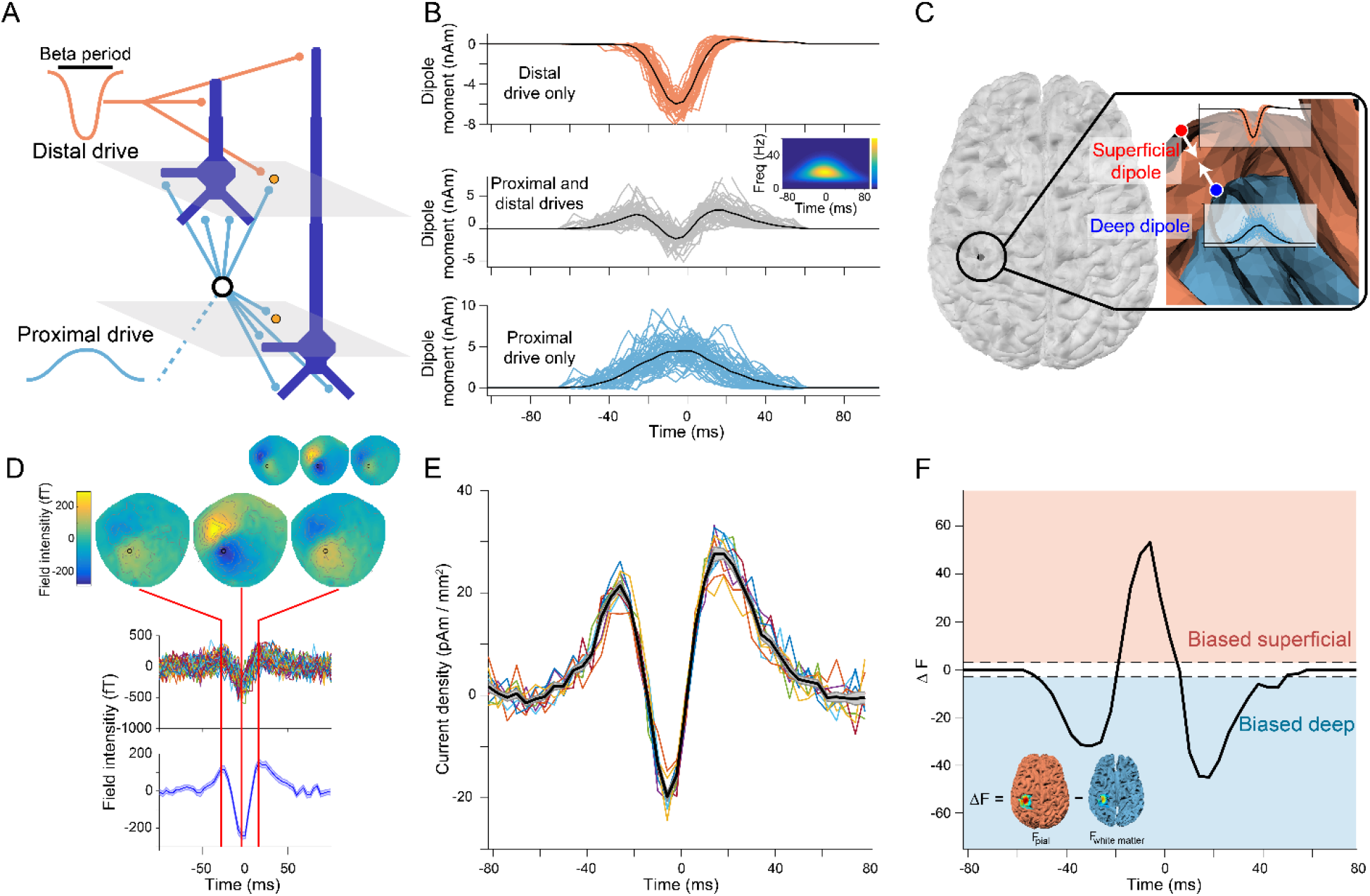
A biophysical model of beta burst generation predicts a bilaminar time course of beta burst dynamics. A) Beta bursts were generated by a model with a broad proximal excitatory synaptic drive temporally aligned with a strong distal synaptic drive. B) The model was run with just the distal drive (top) and just the proximal drive (bottom), and the resulting cumulative dipole moments were used as source signals to generate the simulated MEG sensor data. Each line shows the time series from a single burst simulation (n=50) and the average is shown as a black line. The middle panel shows the cumulative dipole moment generated from the model with both drives, exhibiting the same waveform and timefrequency (inlay) features observed in the human MEG data. C) The generative model included two oppositely oriented currents positioned at corresponding locations on the pial and white matter surfaces in motor cortex, which represent the superficial and deep cortical layers, with source activity given by the model run with the distal and proximal synaptic drives, respectively. D) Simulated sensor data generated by the model has the same spatial and temporal features as beta bursts observed in the human subject MEG data used to determine the simulated current locations and orientations (inlay). *E) Time course of source current density resulting from the localizer source inversion on the simulated sensor datasets. Each simulation (n=10) is shown as a colored line and the black line corresponds to the average over all simulations.* F) The sliding window source inversion correctly identifies that the simulated bursts were generated by activity predominately in deep layers at the beginning and end of the burst, with stronger superficial layer activity at the peak of the burst. The dashed lines (at ΔF = ±3) show the point at which one laminar model is 20 times more likely than the other model.

We then created a generative MEG model based on the anatomy of a human subject and simulated a deep and a superficial source at the location identified by the localizer source inversion for that subject. Both sources were oriented according to the estimated cortical column orientation at that location (Bonaiuto et al., 2020). Following the biophysical model prediction, the superficial current pointed in the direction of the white matter surface (i.e. into the cortex), whereas the deep current flow pointed in exactly the opposite direction (out of the cortex). The time course of each current in the generative MEG model was determined by the cumulative dipole moment of the biophysical model with only the distal (superficial dipole) or proximal (deep layer dipole) synaptic drive applied. This generative model was then used to create simulated MEG sensor data which had the same spatial and temporal features observed in the human MEG data, providing support for the model predictions (**Figure 3**D). This process was repeated 10 times to yield a dataset of virtual subjects. Source inversion on these simulated datasets revealed the same wavelet-like burst waveform shape in source space current density (**Figure 3**E). As expected, the sliding time window source inversion identified the bilaminar pattern of simulated activity used to generate the data (**Figure 3**F): with significantly more evidence for the white matter (deep) model at the beginning and end of the burst (−50 – −18ms and 6 – 50ms) and for the pial (superficial) model around the peak of the burst (−14 – 2ms). These results generalized beyond the particular time series generated by the biophysical model, as we obtained the same results using a highly simplified model with deep and superficial currents given by scaled Gaussian time series (**Figure S1**).

Having validated that the new temporally resolved laminar analysis could correctly reconstruct the time course of the relative strength of deep and superficial layer activity in simulated data, we next wanted to determine if the analysis could distinguish between a range of alternative synthetic generative models which differed in terms of their deep and superficial dipole moment waveforms and polarities. We first tested a synthetic model in which the distal drive consisted of a temporally broad and weak signal, directed into the cortex, whilst the proximal drive consisted of a stronger and briefer signal, directed out of the cortex (**Figure 4**A). This resulted in a simulated sensor dataset with an oppositely oriented field and reversed polarity waveform to that observed in the human subject MEG data (**Figure 4**B). Importantly, the temporally resolved laminar analysis was able to correctly identify that this dataset was generated by a pattern of activity that was stronger in superficial layers at the beginning and end of the burst, and deep layers around the peak of the burst (**Figure 4**C). We next tested a model in which the deep and superficial layer dipoles pointed away rather than towards each other (**Figure 4**D). The simulated sensor data from this model also had oppositely oriented spatial and temporal features compared to those observed in human subject MEG data (**Figure 4**E), and the temporally resolved laminar analysis again correctly identified the time course of laminar activity (**Figure 4**F). We tested a range of other synthetic models including two models containing single deep or superficial dipoles in isolation, and models where dipole currents were oriented in the same direction. In each of these alternative models, the analysis correctly identified the time course of dominant laminar activity (**Figure S2**). The only model whose simulated dataset matched that of the human subject data in terms of field direction and waveform shape was the original biophysical model in which strong superficial layer activity was temporally aligned with broad, weaker activity in deep layers. The superficial layer synaptic activity drove net current flow toward the deep layers and vice versa for the deep layer activity. It should be noted that these alternate synthetic models are not intended to realistically explain the generation of beta bursts in sensorimotor cortex, and the examples in **Figure 4** were generated without using the biophysical model. For example, it is likely biophysically impossible for deep layer synaptic inputs to drive current flow away from the superficial layers. However, these simulations demonstrate that the temporally resolved laminar analysis can, in principle, distinguish between different patterns of laminar activity in human MEG data.

**Figure 4.**
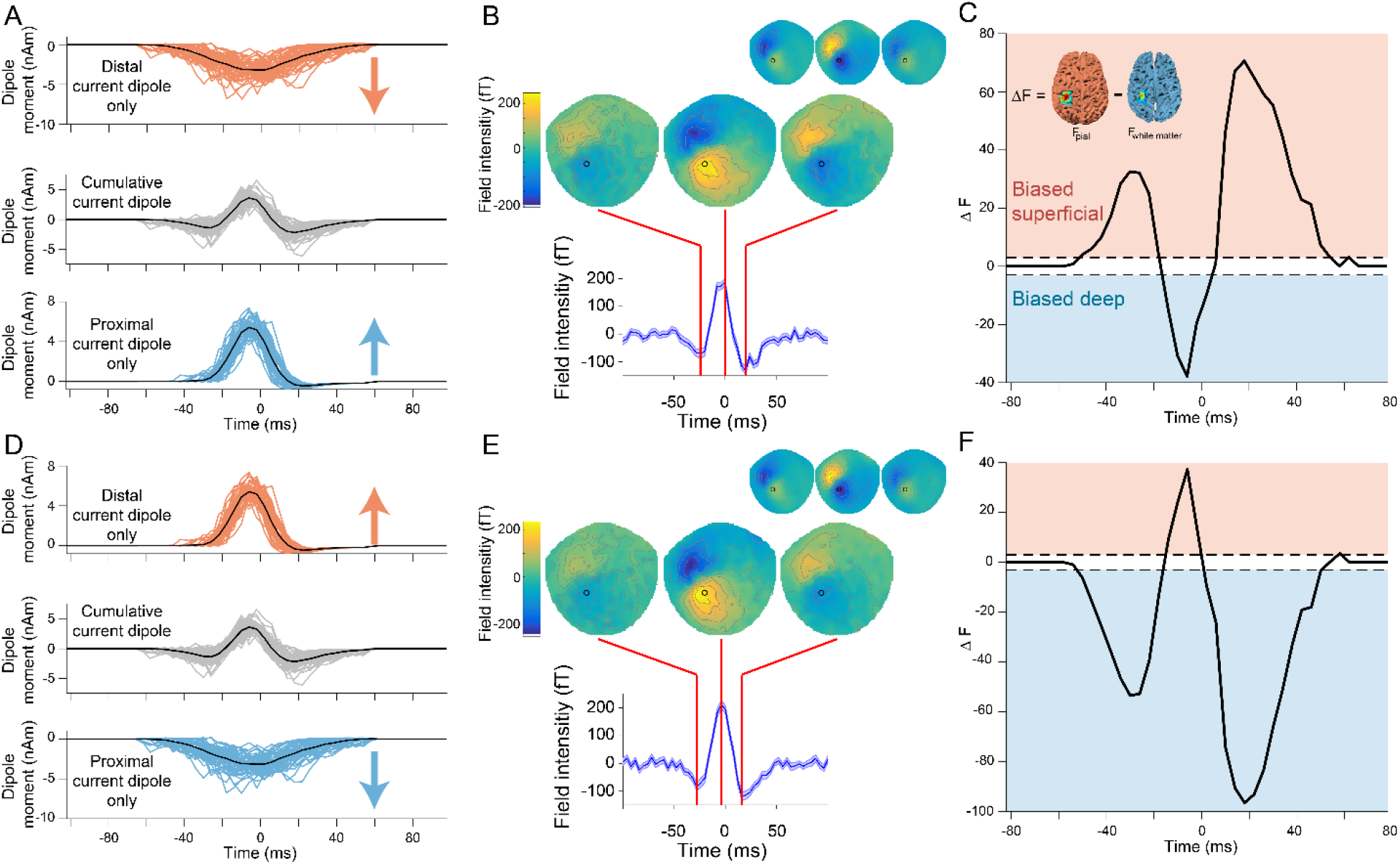
Alternate synthetic models with reversed proximal and distal dipole moment waveforms or polarities generate different predictions. A) Synthetic model with reversed distal (top) and proximal (bottom) dipole moment waveforms from which simulated MEG data were generated. B) The synthetic model with reversed dipole moment waveforms generated simulated sensor data with oppositely oriented spatial and temporal features, compared to beta bursts observed in the human MEG data used to determine the simulated current locations and orientations (inlay). C) The sliding window source inversion correctly identifies that the bursts generated by the reversed dipole moment waveforms model were generated by activity predominately in superficial layers at the beginning and end of the burst, with stronger deep layer activity at the peak of the burst. D) Alternative synthetic model with reversed dipole polarities. E) Similar to the reversed dipole moment waveforms model, the reversed dipole polarity model generated simulated sensor data with oppositely oriented spatial and temporal features, compared to bursts in human MEG data (inlay). F) The sliding time window source inversion can determine the laminar source of dominant activity in the reversed dipole polarity model.

### Temporally resolved estimates of lamina-specific currents require high precision MEG data

The spatial precision of MEG data is classically limited by its signal-to-noise ratio (SNR) and error in localization of the brain relative to the MEG sensors (co-registration error; Hillebrand and Barnes, 2002; Troebinger et al., 2014b). We used subject-specific 3D printed head-casts to reduce within-session movement and reduce between-session head position variability, allowing us to reduce co-registration error and increase SNR by averaging across sessions (Meyer et al., 2017; Troebinger et al., 2014b). Before applying the new analysis to human MEG data, we first wanted to determine if the SNR and co-registration accuracy achieved by this approach were within the range required for accurate estimates of lamina-specific currents. We therefore used the simplified beta burst model (**Figure S1**) to generate simulated MEG datasets with a range of SNR and co-registration error levels, and examined the resulting laminar biases during the tails and peak of the simulated bursts (see Methods). At low SNR levels, no significant laminar biases could be detected, but as SNR increased above −30 dB, the deep bias at the tail ends of the burst and superficial bias at the peak of the burst could be resolved (**Figure 5**A). Moreover, we saw that the ability to resolve this laminar bias slowly decreased with increasing co-registration error, and deep and superficial biases could no longer be reliably detected when this error increased beyond 2 mm and 2 degrees (**Figure 5**B). However, the SNR and co-registration error of our human subject data were well within this bounds (pre-movement burst SNR: M=299.72, SE=2.23 dB; post-movement burst SNR: M=295.15, SE=1.82 dB; between-session movement variability <0.6 mm; within-session movement maximum=0.22 mm; Meyer et al., 2017).

**Figure 5.**
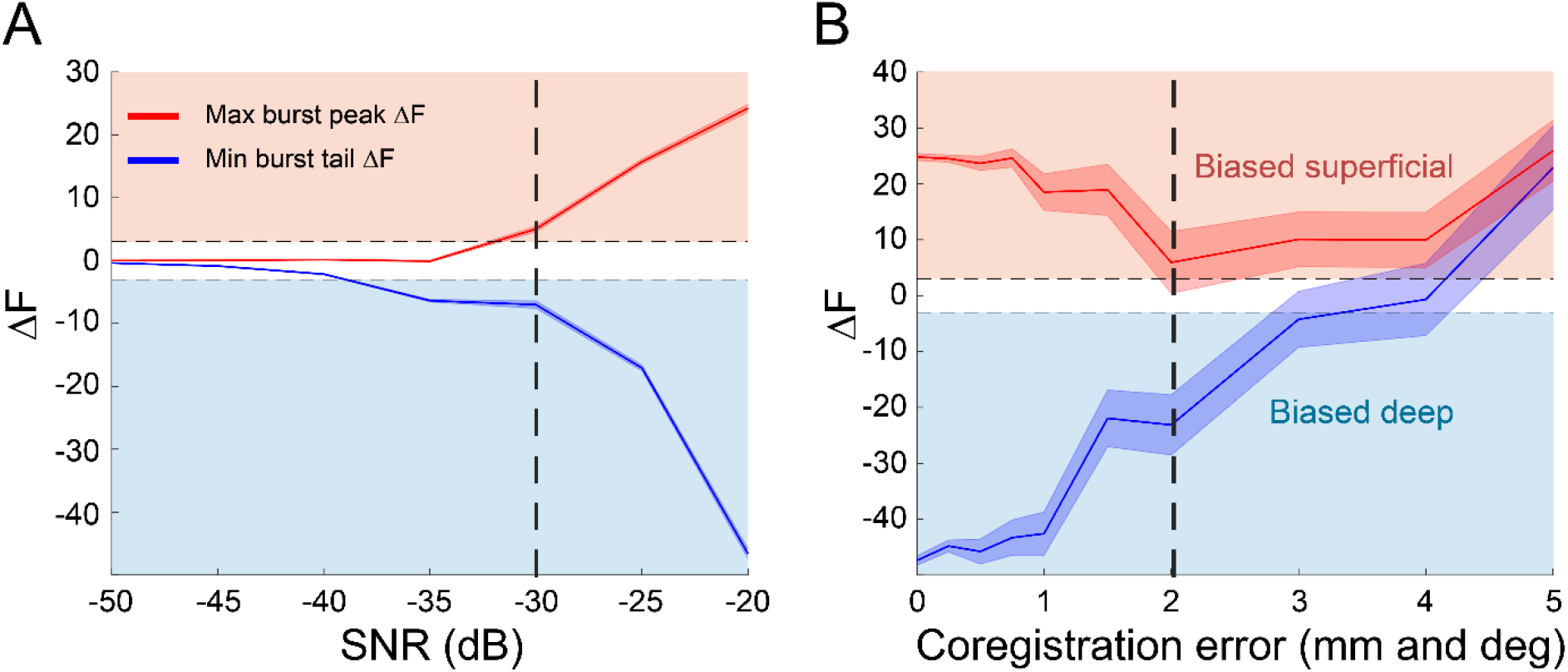
High precision MEG data enables localization of laminar dynamics. A) At SNR levels below −30 dB (dashed vertical line), it is impossible to detect laminar biases during the tail ends and peak of simulated bursts (shaded area shows standard error). Above −30 dB the full time course of laminar activity can be resolved. B) When co-registration error is less than 2 mm and 2 degrees (dashed vertical line), it is possible to resolve the deep and superficial biases during the tails and peak of simulated beta bursts (shaded area shows standard error). Above 2 mm and 2 degrees, laminar bias detection is unreliable.

### The laminar activity pattern of motor beta bursts in human MEG data conforms to predictions from biophysical modelling

After validating the temporally resolved laminar analysis on simulated data, and estimating the required SNR and co-registration accuracy, we then proceeded to apply this analysis to pre- and post-movement beta bursts from human MEG data. Pre-movement beta bursts from motor cortex at the source level had the same wavelet-like current density waveform observed in the sensor data (**Figure 6**A). As predicted by the biophysical model, activity at the beginning and end of pre-movement bursts predominately occurred in deep cortical layers, whereas activity at the peak of the bursts was biased superficially (**Figure 6**B). Prior to the burst peak, the free energy difference reached a minimum of −34.73, indicating that the deep layer model was more strongly supported by the MEG data than the superficial layer model (a free energy difference of ±3 indicates that one model is approximately 20 times more likely than the other). After the burst peak, the minimum free energy difference was −31.55. By contrast, the free energy difference reached a maximum of 49.15 around the peak of burst, suggesting that the superficial layer model was more strongly supported by the data than the deep layer model. Post-movement beta bursts had the same wavelet-like current density waveform shape at the source level (**Figure 6**C), and had the same bilaminar pattern activity as pre-movement bursts and as predicted by the biophysical model (pre-burst minimum ΔF= −71.32; post-burst minimum ΔF= −48.02; burst peak ΔF= 51.47; **Figure 6**D). These results were robust to choice of analysis parameter values (**Figure S3–5**) and individual subject differences (**Figure S6**).

**Figure 6.**
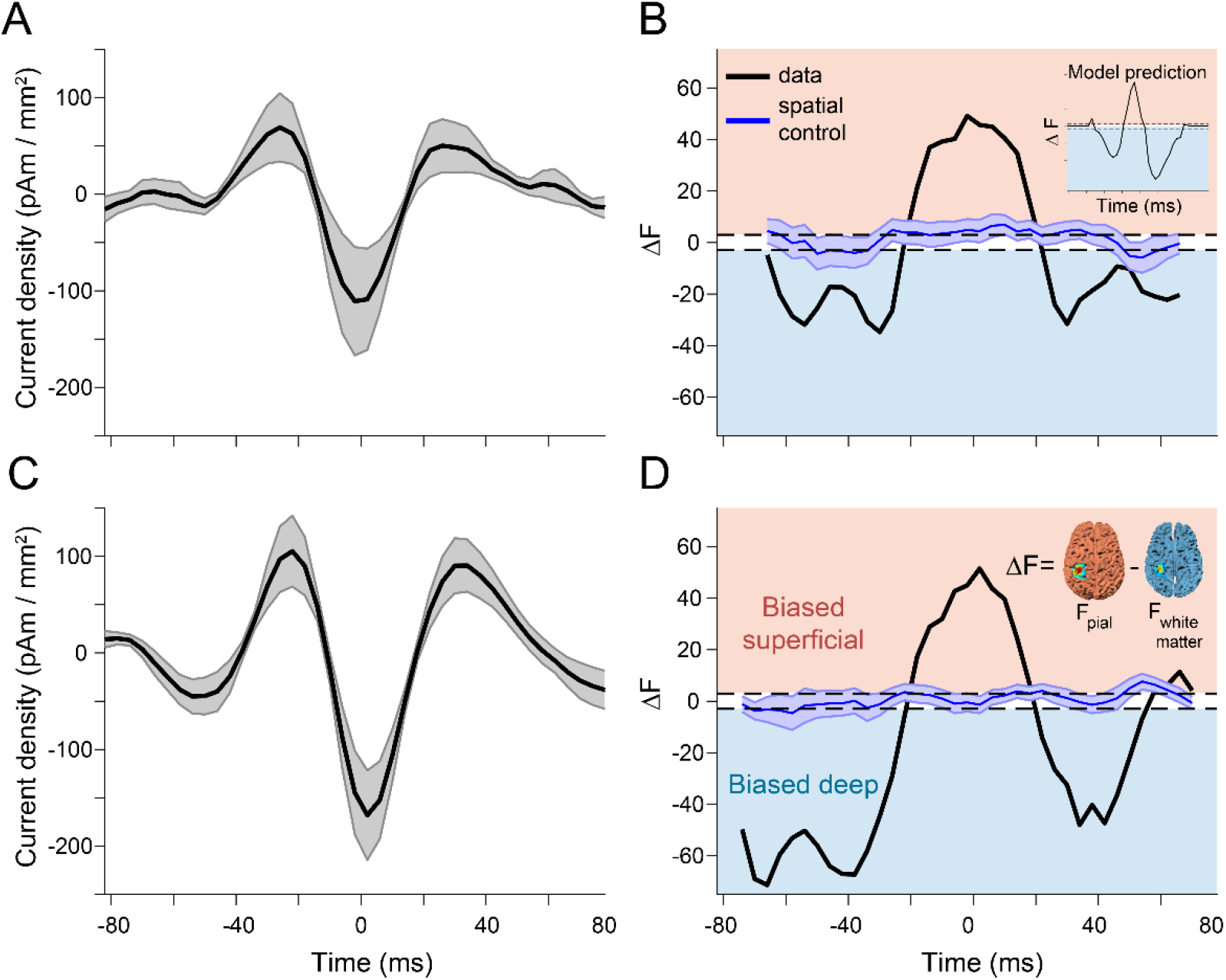
Pre- and post-movement motor beta bursts exhibit the predicted bilaminar dynamics. A) The aligned pre-movement beta burst source level current density time courses averaged across subjects (shaded area shows standard error). B) Activity at the beginning and end of pre-movement bursts localized to deep cortical layers, whereas activity at the peak of the bursts localized superficially. This was not true for spatially randomized (blue) surrogate data which yielded a flat Bayes factor time course that was not biased to either surface (shared area shows standard error). Post-movement beta bursts had similar source level temporal (C) and laminar (D) characteristics as pre-movement bursts.

We ran a control analysis to ensure that these results were spatially specific, using the same burst-averaged data as the main analysis, but with random vertices on the pial surface (and corresponding vertices on the white matter surface) as Bayesian priors in the sliding time window source inversion. This control analysis was designed to ensure that the bilaminar pattern of activity characterizing precentral beta bursts was specific to the cortical location in which these bursts were occurring. The control analysis yielded flat estimates of laminar activity, without any significant bias toward the deep or superficial layers, for both pre- and post-movement bursts (**Figure 6** C,D). This was also true for a temporal control analysis, which involved shuffling the bursts in the time domain before averaging them, prior to running the localizer and sliding time window source inversions (**Figure S7**).

## Discussion

We combined temporally resolved laminar inverse analysis and biophysical modeling to show that movement-related beta bursts in human motor cortex arise from distinct and temporally aligned deep and superficial layer excitatory synaptic drives that generate alternating dominant dipole currents in deep and superficial layers. The deep layer excitatory synaptic input initially drives current toward the superficial layers (i.e., out of the cortex), and this is transiently interrupted by the superficial layer input which drives current in the opposite direction to generate the dominant peak in the beta burst waveform. This pattern of activity is in accordance with predictions from a biophysical model of somatosensory beta burst generation (Jones et al., 2009; Law et al., 2019; Neymotin et al., 2020; Sherman et al., 2016), suggesting that motor and somatosensory beta bursts share a common mechanism.

In addition to identifying the mechanism of beta burst generation in human motor cortex, we validate the use of high precision MEG for temporally resolved estimates of the relative strength of deep and superficial cortical activity. Previously developed techniques were only able to localize the laminar source of induced and temporally averaged activity over a relatively wide time window (Bonaiuto et al., 2018a, 2018b; Troebinger et al., 2014a), and were only able to determine relative laminar biases due to differences in SNR in the beta and gamma band signals analyzed (Bonaiuto et al., 2018a). Here we have extended these methods to resolve the time course of laminar neural dynamics, analyzed putative deep and superficial layer signals with comparable SNR, and taken advantage of the ability of hpMEG to infer the orientation of cortical columns (Bonaiuto et al., 2020) in order to anatomically constrain current orientations and polarity in laminar MEG generative models. The resulting method yields estimates of the time series of laminar dominance reflective of synaptically induced currents that are in-line with depth electrode recordings from rodents and nonhuman primates (Sherman et al., 2016), providing further support for this family of laminar hpMEG analyses.

Motor and sensory cortex differ in terms of their afferent projections from other regions (Arikuni et al., 1988; Felleman and Van Essen, 1991; Friedman et al., 1986; Jones et al., 1978; Tokuno and Tanji, 1993), but have the same layer-specific input patters (Kuramoto et al., 2009). In line with this, our results suggest a common mechanism for the generation of beta bursts across sensorimotor cortex (though this does not imply that all cortico-basal beta bursts are cortically generated). However, beta bursts in these regions have distinct relationships with behavior (Jana et al., 2020; Little et al., 2019; Shin et al., 2017; Wessel, 2020), for example, pre- and post-movement bursts in motor cortex are related to different aspects of motor responses (Little et al., 2019). Identifying the source of the deep and superficial layer projections to motor cortex that drive pre- and post-movement beta bursts will help to shed light on their functional role. Future extensions to the technique developed here could identify these sources by incorporating measures of time-varying functional connectivity (Astolfi et al., 2008; Hesse et al., 2003; Youssofzadeh et al., 2016), allowing for inference of lamina-specific connectivity.

There are some limitations to the temporally resolved laminar analysis presented here. Like previously developed laminar hpMEG techniques (Bonaiuto et al., 2018a, 2018b), the new analysis can only determine the relative bias of deep versus superficial layer activity; it cannot recover the time course of activity in each layer. However, coupling the analysis with a biophysical model and a range of alternative synthetic models allows inference of lamina-specific activity time courses by determining the directionality of deep and superficial currents. Future extensions involving generative models with surfaces representing intermediate cortical layers, and higher SNR data afforded by cryogen-free MEG sensors (optically-pumped magnetometers; OPMs; Boto et al., 2018, 2016; Holmes et al., 2018; Iivanainen et al., 2019, 2017; Knappe et al., 2014), may be able to use an approach similar to current source density methods (Pettersen et al., 2008; Rappelsberger et al., 1981; Schroeder et al., 1998, 1991, 1990) to increase the spatial specificity of laminar inference.

Although we found deep and superficial biases at different periods over the time course of beta bursts, these results are in fact consistent with the finding that temporally averaged pre- and post-movement induced beta activity predominates in deep cortical layers (Bonaiuto et al., 2018a). The biophysical model adopted here generates beta bursts consisting of two tails of deep layer activity and a transient peak of superficial layer activity, resulting in an overall bias towards deep layers. We compensated for this predicted net deep layer bias by using a pial surface localizer, but future work could overcome potential depth biases by adopting an approach using localization with an intermediate layer surface or temporal basis functions generated by biophysical model predictions. More generally, these results question the notion that activity in various frequency channels is simply segregated into different cortical layers. Instead, they suggest a view in which the generation of frequency-specific activity has a temporal and laminar profile, and only appears to be restricted to deep or superficial layers when measuring the net bias through the lens of temporally averaging induced activity.

Another potential confound is patch size. In earlier work (Troebinger et al., 2014a) we showed that underestimation of the true patch extent will tend to bias estimates towards deeper layers, whereas an overestimation of patch extent will tend to bias layer estimates more superficially. In our parameter sensitivity analyses (**Figure S4, S5**), however, we found laminar inference to be robust over a range of realistic patch sizes. Here we had a specific assumption that activity is driven from a combination of deep and superficial sources with equal patch extents. However, we should mention that an alternative hypothesis, which we did not explore, is that the bursts arise from a single layer within an expanding and contracting local coherence in current flow.

Our analysis aligned MEG sensor data to beta amplitude peaks and treated these signals as event-related fields (ERFs). Sensory and motor ERFs (and event-related potentials; ERPs) have waveforms which are composed of a series of dynamic components generated by temporal patterns of activity in both deep and superficial cortical layers (Mehta et al., 2000a, 2000b; Olson, 2001; Schall et al., 2020; Schroeder et al., 1992, 1991; Szymanski et al., 2011), but studies combining laminar electrode and ERP recordings from nonhuman primates are rare (Woodman, 2012). The sliding time window inversion technique presented here can also be used to non-invasively test hypotheses, potentially constrained by biophysical modeling (Jones et al., 2009, 2007; Neymotin et al., 2020), concerning the generation of event-related neural activity by interlaminar dynamics.

The finding that temporally transient deep and superficial layer excitatory synaptic drives generate movement-related beta bursts in human motor cortex provides relevant mechanistic constraints to theories of the functional relevance of pre- and post-movement beta activity (Alegre et al., 2008, 2002; Cao and Hu, 2016; Cassim et al., 2001; Classen et al., 1998; Engel and Fries, 2010; Fetz, 2013; Jasper and Penfield, 1949; Murthy and Fetz, 1992; Pfurtscheller et al., 1996; Reyns et al., 2008; Roelfsema et al., 1997; Saleh et al., 2010; Spitzer and Haegens, 2017; Tan et al., 2016, 2014). Such theories are often framed in terms of slowly modulated beta oscillations, and generally predict sustained beta activity arising from either sustained inputs or recurrently maintained activity. On the contrary, the transitory confluence of deep and superficial layer synaptic drives in beta burst generation suggests a dynamic role for motor beta bursts (Little et al., 2019), such as the integration of different sensorimotor information signals during pre- and post-movement periods.

## Methods

Data from eight healthy, right-handed, volunteers with normal or corrected-to-normal vision and no history of neurological or psychiatric disorders were used for our analyses (Bonaiuto et al., 2018a; Little et al., 2019). The study protocol was in accordance with the Declaration of Helsinki, and all participants gave written informed consent which was approved by the UCL Research Ethics Committee (reference number 5833/001). All analysis code is available at http://github.com/jbonaiuto/beta_burst_layers.

### MRI acquisition

Two MRI scans were acquired prior to MEG scanning with a 3T whole body MR system (Magnetom TIM Trio, Siemens Healthcare, Erlangen, Germany) using the body coil for radio-frequency (RF) transmission and a standard 32-channel RF head coil for reception. The first was a standard T1-weighted scan for creation of each subject’s individual head-cast (Meyer et al., 2017), and the second was a high resolution, quantitative multiple parameter map (MPM; Weiskopf et al., 2013) for MEG source location.

The T1-weighted protocol used a 3D spoiled fast low angle shot (FLASH) sequence with 1 mm isotropic image resolution, field-of view of 256, 256, and 192 mm along the phase (anterior-posterior, A–P), read (head-foot, H–F), and partition (right-left, R–L) directions, respectively, repetition time of 7.96 ms, and excitation flip angle set to 12°. A single echo was acquired after each excitation to yield a single anatomical image. A high readout bandwidth (425 Hz/pixel) was used to preserve brain morphology, with no significant geometric distortions observed in the images. The acquisition time for this sequence was 3min 42s. A 12-channel head coil was used for signal reception without either padding or headphones in order to avoid introduction of scalp distortions.

The MPM protocol was comprised of three differentially-weighted, RF and gradient spoiled, multi-echo 3D fast low angle shot (FLASH) sequences and two additional calibration sequences to correct for inhomogeneities in the RF transmit field (Callaghan et al., 2015; Lutti et al., 2012, 2010), with whole-brain coverage at 800 μm isotropic resolution.

The FLASH sequences had predominantly proton density (PD), T1, or magnetization transfer saturation (MT) weighting. The PD- and MT-weighted volumes used a flip angle of 6°, and the T1 weighted acquisition used a flip angle of 21°. MT-weighting was achieved through the application of a Gaussian RF pulse 2 kHz off resonance with 4 ms duration and a nominal flip angle of 220° prior to each excitation. The field of view was 256 mm head-foot, 224 mm anterior-posterior (AP), and 179 mm right-left (RL). Gradient echoes were acquired with alternating readout gradient polarity at eight equidistant echo times ranging from 2.34 to 18.44 ms in steps of 2.30 ms using a readout bandwidth of 488 Hz/pixel, but only six echoes were acquired for the MT-weighted volume in order to maintain a repetition time (TR) of 25 ms for all FLASH volumes. Partially parallel imaging using the GRAPPA algorithm was employed to accelerate data acquisition, with a speed-up factor of 2 in each phase-encoded direction (AP and RL) with forty integrated reference lines.

To maximize the accuracy of the measurements, inhomogeneity in the transmit field was mapped by acquiring spin echoes and stimulated echoes across a range of nominal flip angles following the approach described in Lutti et al. (2010), including correcting for geometric distortions of the EPI data due to B0 field inhomogeneity. The total acquisition time for all MRI scans was less than 30 min.

Quantitative maps of proton density (PD), longitudinal relaxation rate (R1 = 1/T1), MT and effective transverse relaxation rate (R2* = 1/T2*) were subsequently calculated (Weiskopf et al., 2013).

### FreeSurfer surface extraction

FreeSurfer (v5.3.0; Fischl, 2012) was used to reconstruct pial and white matter surfaces from the MPM volumes for source localization of MEG sensor data. We used a custom FreeSurfer procedure to process MPM volumes, using the PD and T1 volumes as inputs (Carey et al., 2017), resulting in surface meshes representing the pial surface (adjacent to the cerebro-spinal fluid, CSF), and the white/grey matter boundary. We used a custom routine to downsample each surface by a factor of 10 while maintaining the correspondence between surface vertices (Bonaiuto et al., 2020). This yielded two meshes of the same size (same number of vertices and edges), containing about 30,000 vertices each (M = 30,094.8, SD = 2,665.5 over participants).

### Head-cast construction

From an MRI-extracted 3D scalp model, we constructed a head-cast that fit between the participant’s scalp and the MEG dewar (Bonaiuto et al., 2018a; Meyer et al., 2017; Troebinger et al., 2014b). Scalp surfaces were first extracted from the T1-weighted volumes acquired in the first MRI protocol using SPM12 (http://www.fil.ion.ucl.ac.uk/spm/). This tessellated surface, along with 3D models of fiducial coils placed on the nasion and the left and right pre-auricular points, was then placed inside a virtual version of the scanner dewar in order to minimize the distance to the sensors while ensuring that the participant’s vision was not obstructed. The resulting model (including spacing elements and ficudial coil protrusions) was printed using a Zcorp 3D printer (Zprinter 510). The 3D printed model was then placed inside a replica of the MEG dewar and polyurethane foam was poured in between the model and dewar replica to create the participant-specific head-cast. The fiducial coil protrusions in the 3D model therefore became indentations in the foam head-cast, into which the fiducial coils were placed during scanning. The locations of anatomical landmarks used for co-registration are thus consistent over repeated scans, allowing us to merge data from multiple sessions (Bonaiuto et al., 2018a; Meyer et al., 2017).

### Behavioral task

Participants completed a visually cued, action-based decision making task in which they responded to an instruction cue projected on a screen by pressing one of two buttons using the index and middle finger of their right hand (Bonaiuto et al., 2018a). After a baseline fixation period, a random dot kinematogram (RDK) was displayed for 2s with coherent motion to the left or to the right. Following a 500ms delay period, an instruction cue (an arrow pointing to the left or the right), prompted participants to respond by pressing either the left or right button. The level of RDK motion coherence and the congruence between the RDK motion direction and instruction cue varied from trial to trial, but for the purposes of the present study, we collapsed across conditions and analyzed beta activity before and after all button press responses. See Bonaiuto et al. (2018a) for a complete description of the task paradigm and structure.

Each block contained a total of 180 trials. Participants completed three blocks per session, and 1–5 sessions on different days, resulting in 540–2700 trials per participant (M = 1822.5, SD = 813.2). The task was implemented in MATLAB (The MathWorks, Inc., Natick, MA) using the Cogent 2000 toolbox (http://www.vislab.ucl.ac.uk/cogent.php).

### MEG acquisition and preprocessing

MEG data were acquired using a 275-channel Canadian Thin Films (CTF) MEG system with superconducting quantum interference device (SQUID)-based axial gradiometers (VSM MedTech, Vancouver, Canada) in a magnetically shielded room. A projector was used to display visual stimuli on a screen (~50 cm from the participant), and a button box was used for participant responses. The data collected were digitized continuously at a sampling rate of 1200 Hz. MEG data preprocessing and analyses were performed using SPM12 (http://www.fil.ion.ucl.ac.uk/spm/) using MATLAB R2014a. The data were filtered (5th order Butterworth bandpass filter: 2–100 Hz, Notch filter: 50 Hz) and downsampled to 250Hz. Eye blink artifacts were removed using multiple source eye correction (Berg and Scherg, 1994). Trials were then epoched from 2s before the participant’s response to 2s after. Blocks within each session were merged, and trials whose variance exceeded 2.5 standard deviations from the mean were excluded from analysis. Preprocessing code is available at http://github.com/jbonaiuto/meg-laminar.

### Burst definition and analysis

Burst were defined used a two-stage process. Firstly, the sensor data time series was inverted onto the subject specific pial surface mesh (note the explicit bias here towards the pial surface is simply to obtain a robust time series estimate so that bursts can be identified) using an Empirical Bayesian beamformer algorithm (EBB; Belardinelli et al., 2012; López et al., 2014) as implemented in SPM12. Source inversion was completed for a 5 second window of data epoched to the movement cue for the pre-movement cue and to a 4 second window of data epoched to the button press for the post-movement cue. These data were projected into 274 orthogonal spatial (lead field) modes and 4 temporal modes and current density time series were created by multiplying the sensor data by the weighting matrix (M) between sensors and source from the inversion, and the data reduction matrix (U) that specifies the significance of the data modes that map to the cortex. The time series from the vertex closest to the center of the hand knob in the primary motor cortex was then taken forward for burst analysis. The selected current density time series was filtered using a 4^th^ order Butterworth filter in the beta band (13 – 30 Hz) and the amplitude determined using the Hilbert function. Bursts were defined using an empirically defined threshold (Little et al., 2019) of 1.75 standard deviations above the median beta amplitude. Sensor data was then re-epoched around the peak of the beta burst amplitude (+-100ms). Additionally, a more accurate epoching of bursts was achieved through using the time series data and a Woody filter (Woody, 1967) to align each burst epoch to the average burst template (equivalent to cross correlating individual bursts to the average burst and shifting the epoch window according to the lag that maximizes the correlation). This new burst aligned sensor dataset was then used for the time-resolved laminar analyses.

Burst amplitude was compared between pre- and post-movement epochs using a linear mixed model in R (v3.6.3; R Core Team, 2020) with the lme4 package (v1.1-21; Bates et al., 2014). Minimum beta amplitude in the 50ms surrounding the burst peak was treated as the dependent measure, with epoch as a fixed effect (pre- or post-movement), and subject-specific intercepts as a random effect. Fixed effect significance was tested using a Type II Wald chi-square test.

### Localizer source reconstruction

Source inversion for beta burst localization was performed using the empirical Bayesian beamformer algorithm (EBB) as implemented in SPM12. The source inversion was applied to a 200ms time window, centered on the peak of the average beta burst time course, without Hann windowing. These data were projected into 274 orthogonal spatial (lead field) modes and 4 temporal modes. These inversions used a spatial coherence prior (Friston et al., 2008) with a FWHM of 5 mm. We used the Nolte single shell head model (Nolte, 2003) with the source locations constrained by the vertices of the downsampled cortical surface, and orientations constrained according to the link vectors between the pial and white matter surfaces to approximate the orientation of cortical columns (Bonaiuto et al., 2020). Clusters of vertices with activity above a threshold of 80% of the maximum activity, within a mask of 50mm centered on the hand knob of the left precentral gyrus were carried forward to the sliding time window source reconstruction.

Induced pre- and post-movement beta activity localizes primarily to deep cortical layers (Bonaiuto et al., 2018a). This is also predicted by the biophysical model, which generates beta bursts consisting of two tails reflecting activity in deep layers and one brief peak of superficial layer activity, resulting in a net bias towards deep layers. We confirmed this by performing a Bayes factor comparison between EBB source inversions using pial versus white matter generative models over the entire burst time course, which yielded greater model evidence for the white matter generative model. To account for this predicted net laminar bias and increase sensitivity to potential superficial layer activity, we used the pial surface to localize beta bursts for the subsequent sliding time window source reconstruction.

### Sliding time window source reconstruction

We used an adaptive Woody filter to align the beta burst source time series across subjects (Woody, 1967). All subsequent analysis was based on these temporally aligned datasets. The sliding time window source reconstruction was performed using the Multiple Sparse Priors (MSP; Friston et al., 2008) algorithm as implemented in SPM12. This was applied to a 40ms time window of the aligned average beta burst time course, with Hann windowing, with a frequency of interest of 1-256Hz. As with the localizer source reconstruction, we used a spatial coherence prior with a FWHM of 5mm, and the Nolte single shell model (Nolte, 2003) with link vector orientations (Bonaiuto et al., 2020). The time window was advanced along the time course of the average beta burst in increments of one time step (4ms), and the Bayesian model evidence (approximated by free energy) for the generative model was computed within each window. Within each cluster of vertices identified by the localizer source reconstruction, the sliding time window source reconstruction was conducted using all vertices with a link vector angle within 0.1 radians of that of the cluster peak vertex (Bonaiuto et al., 2020). For each selected vertex, the sliding time window MSP inversion was conducted using the pial surface to constrain source locations with the prior set as the cluster vertex, and again using the white matter surface with the prior set as the corresponding vertex on the white matter surface. The difference in the free energy time series between these inversions (ΔF = F_pial_ – F_white matter_) was then averaged over vertices within each cluster, and then across clusters. These free energy difference time series of the aligned burst data were then summed over subjects to yield a fixed effects estimate of laminar dominance dynamics.

### Biophysical model

We used the open-source Human Neocortical Neurosolver (HNN) software to simulate our biophysical model of a local neocortical microcircuit under exogenous layer specific synaptic drive (https://hnn.brown.edu; Neymotin et al., 2020). HNN’s model, and the beta burst mechanism, was fully described and validated in prior publications (Law et al., 2019; Sherman et al., 2016; Shin et al., 2017). HNN’s underlying neural model simulates the primary electrical currents in the neocortex that create EEG/MEG signals. The model simulations are based on the biophysical origins of the primary electrical currents (i.e., current dipoles), assumed to be generated by the post-synaptic, intracellular longitudinal current flow in the long and spatially aligned dendrites of a large population of synchronously activated neocortical pyramidal neurons. HNN simulates the primary currents from a canonical model of a layered neocortical column via the net intracellular electrical current flow in the pyramidal neuron dendrites, in the direction aligned with the apical dendrites, multiplied by their length (units nano-Ampere-meters). A scaling factor is applied to the net current dipole output to match the amplitude of recorded data and provides an estimate of the number of neurons contributing to the recorded signal. Simulated multicompartment pyramidal (PN) and single compartment interneurons (IN) are arranged in supra- and infra-granular layers, as shown in **Figure 3**A. Neurons receive excitatory synaptic input from simulated trains of action potentials in predefined temporal profiles (e.g. see schematic Gaussians in **Figure 3**A) that activate excitatory synapses at the location of the *proximal* apical/basal and *distal* apical dendrites of the pyramidal neurons, representing activation from thalamic core and thalamic matrix/corticocortical feedback, respectively.

In the simulations described in this paper, we used a modified version of the default parameter set distributed with HNN that simulates beta bursts, namely AlphaAndBeta.param file. In brief, this simulation contained 100PN and 35IN per layer and used a stochastic sequence of 10 Hz proximal excitatory synaptic drive, simultaneous with distal 10 Hz excitatory synaptic drive. The histogram of driving spikes on each cycle of the rhythmic inputs that generated this drive had a Gaussian profile comparable to that shown schematically in Figure 3A. We have shown that this pattern of input can generate bursts of activity with the characteristic beta event waveform shape that emerges as part of the more continuous somatosensory mu-rhythm (Jones et al., 2009; Sherman et al., 2016). Here, we examine the model output corresponding to individual bursts only. Individual beta bursts or “events” occur on cycles of the stochastic 10Hz drive when broad (~100ms) upward current flow from proximal inputs is synchronously disrupted by downward stronger and faster (~50ms) current flow from distal inputs, where the ~50ms duration of the distal drive creates the dominant beta peak and sets the frequency of the oscillation (Sherman et al., 2016). We used three sets of simulations in order to ascertain the contribution of each cortical layer to the beta waveform shape: 1) proximal drive only (**Figure 3**B top), 2) distal drive only (**Figure 3**B bottom); and 3) combined proximal and distal inputs (**Figure 3**B middle). Each simulation type was run across 50 trials, where a trial (170ms) refers to a single execution of the model with a defined set of simulation parameters. Results varied across trials with identical parameters due to the stochastic nature of the exogenous proximal and distal drives: on each cycle of the 10Hz drive, the timing of the synaptic drive is chosen from a Gaussian distribution with mean inter-cycle input time of 100 ms ± 20 ms standard deviation for proximal drive and 100 ms ± 7 ms for distal drive. The mean delay between the proximal and distal drive is 0ms. Note that the distal drive synaptic weights provided to supra- and infra-granular pyramidal neurons were increased from HNN’s default value to 6e-5 μS. The distal and proximal current dipole moments shown in **Figure 3**B were normalized by the maximum of the averaged amplitudes of each, then scaled to match the amplitudes of recorded data. All source code is provided online at https://hnn.brown.edu and https://github.com/jonescompneurolab/hnn, and a parameter file specific to the simulations here will be distributed upon publication.

### Generation of simulated MEG datasets

All simulations were based on a dataset acquired from one human participant, which was used to determine the sensor layout, sampling rate (1200 Hz, downsampled to 250 Hz), number of samples (51), and simulated dipole locations and orientations for the simulations. In each simulation, we specified spatially distributed source activity centered at a single vertex on the pial surface and the corresponding vertex on the white matter surface, approximating the deep and superficial ends of a cortical column (Bonaiuto et al., 2020). The orientations of the simulated dipoles were defined by the vector connecting the two vertices (link vector) with opposite polarities: the pial dipole pointed toward the white matter dipole and vice versa. The time course of simulated activity at the pial vertex was given by the cumulative dipole moment from 50 beta bursts generated by the biophysical model run with only the distal drive applied, and that of the white matter vertex by the cumulative dipole moment from 50 bursts generated by the model run with only the proximal drive applied. The amplitudes of these time courses were scaled to yield beta bursts matching those seen in the human data, with the pial and white matter vertex dipoles having mean magnitudes of 8 and 6 nAm, respectively. We then spatially smoothed these simulated dipole time courses with a Gaussian kernel (FWHM = 5 mm), to obtain two patches of spatially distributed activity. We then used a single shell forward model (Nolte, 2003) to generate a synthetic dataset from the simulated source activity. Unless otherwise specified, Gaussian white noise was added to the simulated data and scaled in order to yield a per-burst amplitude SNR level (averaged over all sensors) of −20 dB.

Simulated data from the simplified synthetic model (**Figure S1**) were generated in the same way as with the biophysical model, but the pial and white matter vertex dipoles had Gaussian activity time courses, centered within the epoch. The pial vertex Gaussian had a width of 10 ms and a magnitude of 6 nAm, and the white matter vertex Gaussian had a width of 25 ms and a magnitude of 4.5 nAm.

### SNR and co-registration error simulations

Beta bursts were generated using the simplified model and Gaussian white noise was added to the simulated data and scaled in order to yield per-trial amplitude SNR levels (averaged over all sensors) between −50 and −20 dB to generate synthetic datasets across a range of realistic SNRs (typically ranging from −40 to −20dB; Goldenholz et al., 2009). This was repeated 50 times per SNR level and the minimum free energy difference during the tails of the burst (−40 – −10 ms and 10 – 40 ms) and maximum free energy difference during the burst peak (−10 – 10 ms) was computed.

Similarly, we simulated between-session co-registration error by introducing a linear transformation of the fiducial coil locations in random directions (0 mm translation and 0 degrees rotation up to 2 mm translation and 2 degrees rotation) to generate a realistic range of uncertainty concerning the location of the brain relative to the MEG sensors (Adjamian et al., 2004; Gross et al., 2013; Ross et al., 2011; Singh et al., 1997; Stolk et al., 2013; Whalen et al., 2008). In these simulations, the per-trial amplitude SNR was set to −20 dB. This was repeated 50 times per co-registration error level, and the minimum free energy difference during the tails of the burst and maximum free energy difference during the burst peak was computed.

## Acknowledgements

JB was supported by the European Research Council (ERC) under the European Union’s Horizon 2020 research and innovation programme (ERC consolidator grant 864550 to J Bonaiuto). SL was funded by a clinical research training grant from the Wellcome Trust (105804/Z/14/Z). SAN was funded by an NIDCD (R01DC012947-06A1) and an Army Research Office Grant (W911NF-19-1-0402). SRJ and SAN were supported by NIH R01EB022889 and NIH R01MH106174.The Wellcome Centre for Human Neuroimaging is supported by a strategic award from Wellcome (091593/Z/10/Z). The funders had no role in the preparation of the manuscript.

## Supplemental Figures

**Figure S1.**
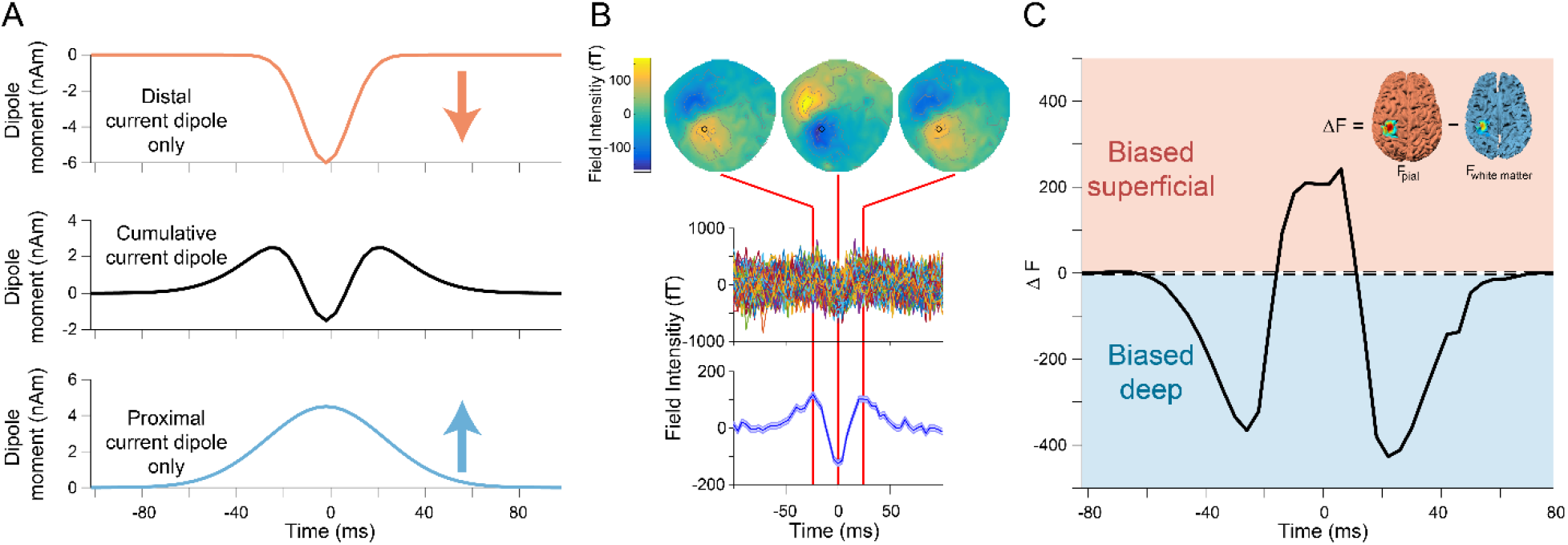
A simple model of beta burst generation yields the same bilaminar predictions as the biophysical model. A) In the simple model, the proximal and distal drives were modeled as Gaussian signals at oppositely oriented dipoles positioned at corresponding locations on the pial (top) and white matter (bottom) surfaces. The resulting cumulative dipole moment exchibited the same waveform features generated by the biophysical model and observed in the human MEG data (middle). B) Simulated sensor-data generated by the model has the same spatial and temporal features as beta bursts generated by the biophysical model and observed in human MEG data. C) The sliding window source inversion correctly identifies that the simulated bursts were generated by activity predominately in deep layers at the beginning and end of the burst, and predominately superficial layer activity at the peak of the burst.

**Figure S2.**
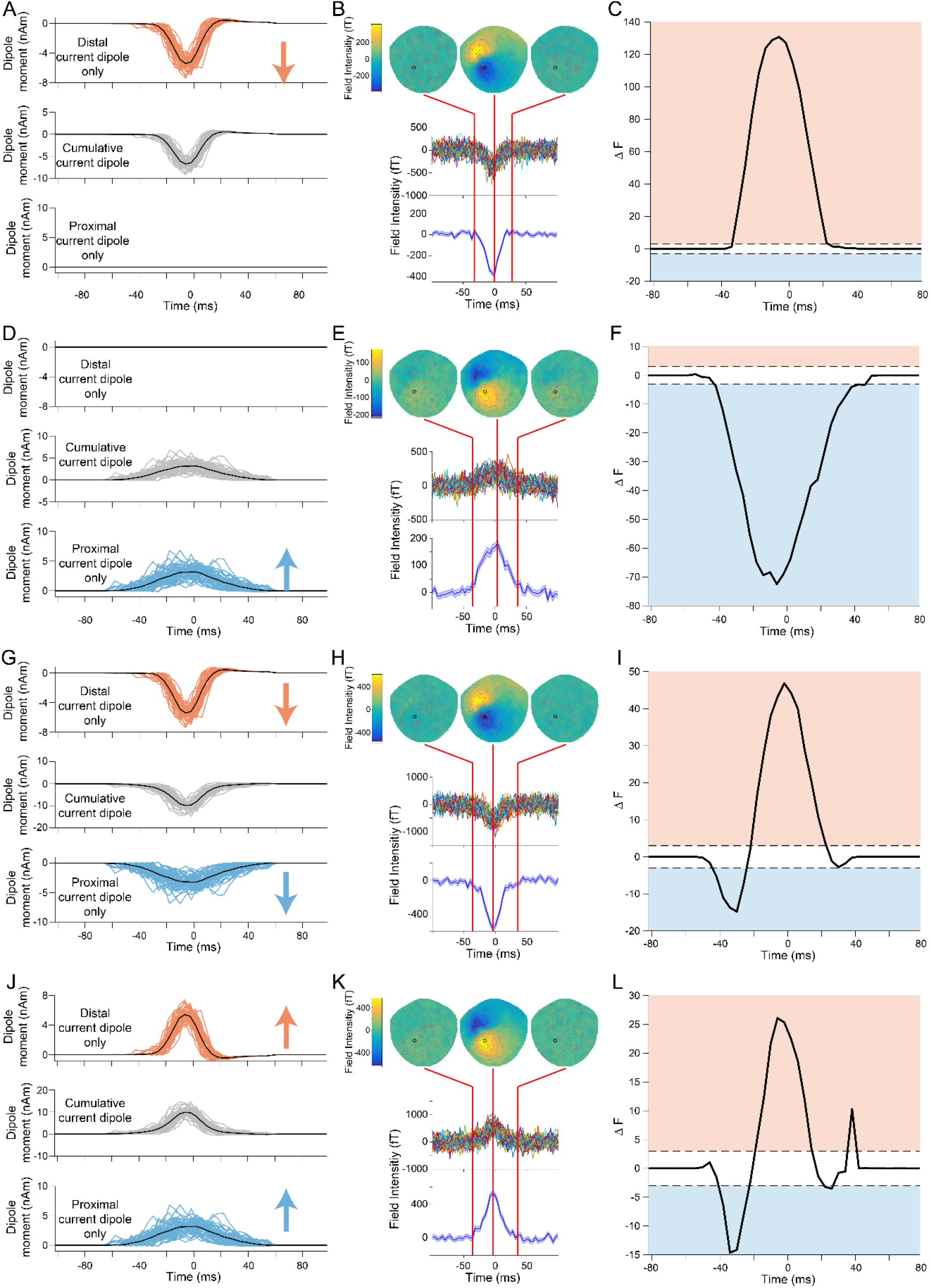
Sliding time window source inversion can correctly identify the laminar time course of alternate synthetic models. Alternate synthetic models including a single superficial dipole (A-C), a single deep layer dipole (D-F), a deep and superficial dipole with oriented in the direction of the deep surface (G-I), or a deep and superficial dipole oriented in the direction of the superficial surface (J-L). The sliding time window inversion correctly predicted superficial layer activity (C) and deep layer activity (F) for the single dipole models, and predominately superficial layer activity with small deep layer biases at the beginning and end of the burst for single orientation direction models (I, L).

**Figure S3.**
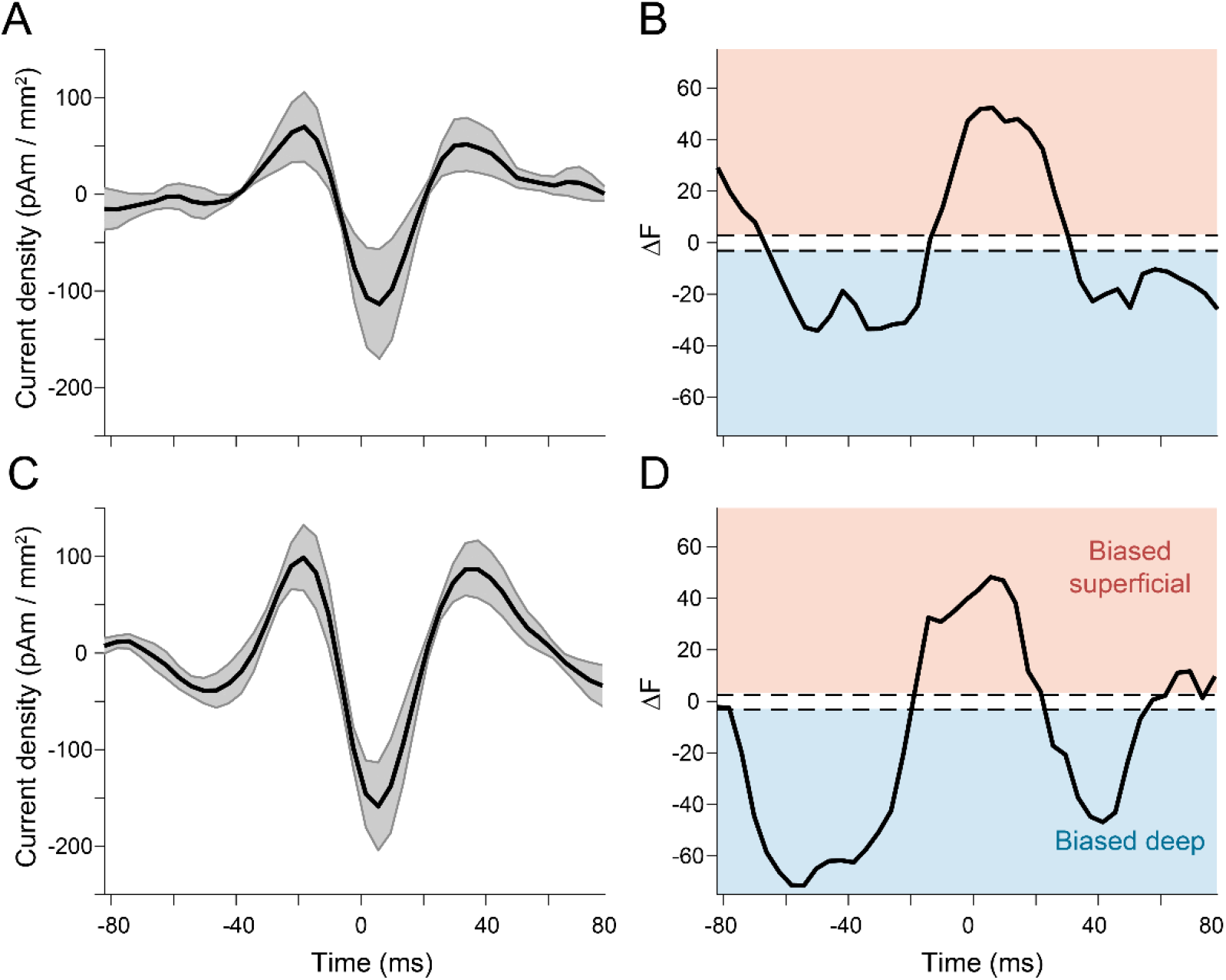
Unaligned beta bursts yield the same results as aligned bursts. A) Unaligned pre-movement beta burst source level current density time courses averaged across subjects (shaded area shows standard error). B) As with the aligned pre-movement bursts (Figure 6), activity at the beginning and end of pre-movement bursts localized to deep cortical layers, whereas activity at the peak of the bursts localized superficially. This was also true for unaligned post-movement beta bursts (C, D).

**Figure S4.**
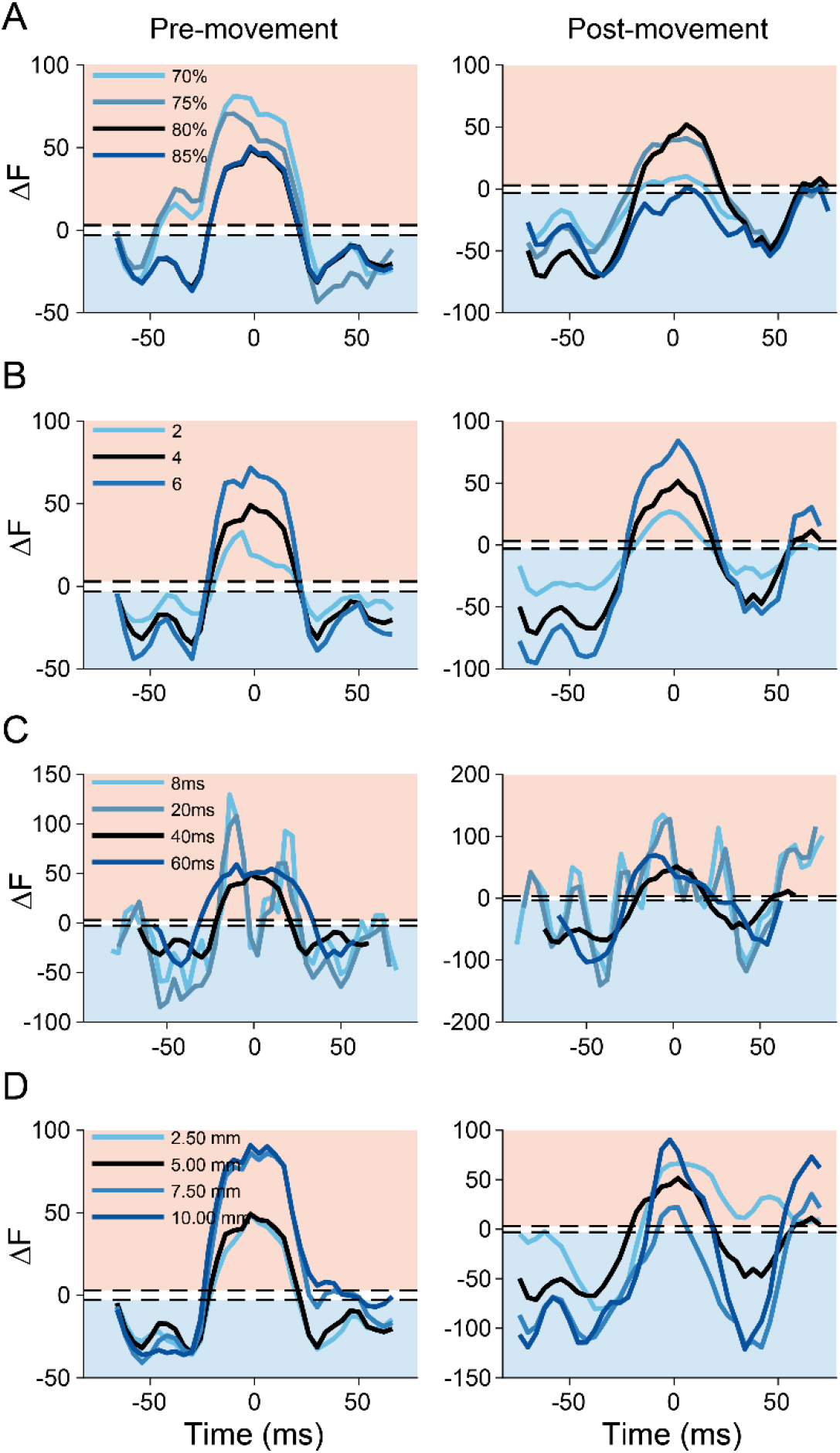
Results with aligned bursts are robust to analysis parameters. Results for aligned pre-movement (left column) and post-movement bursts (right column) were similar across a range of values for cluster thresholding (A), number of temporal models (B), sliding window width (C), and patch size (D).

**Figure S5.**
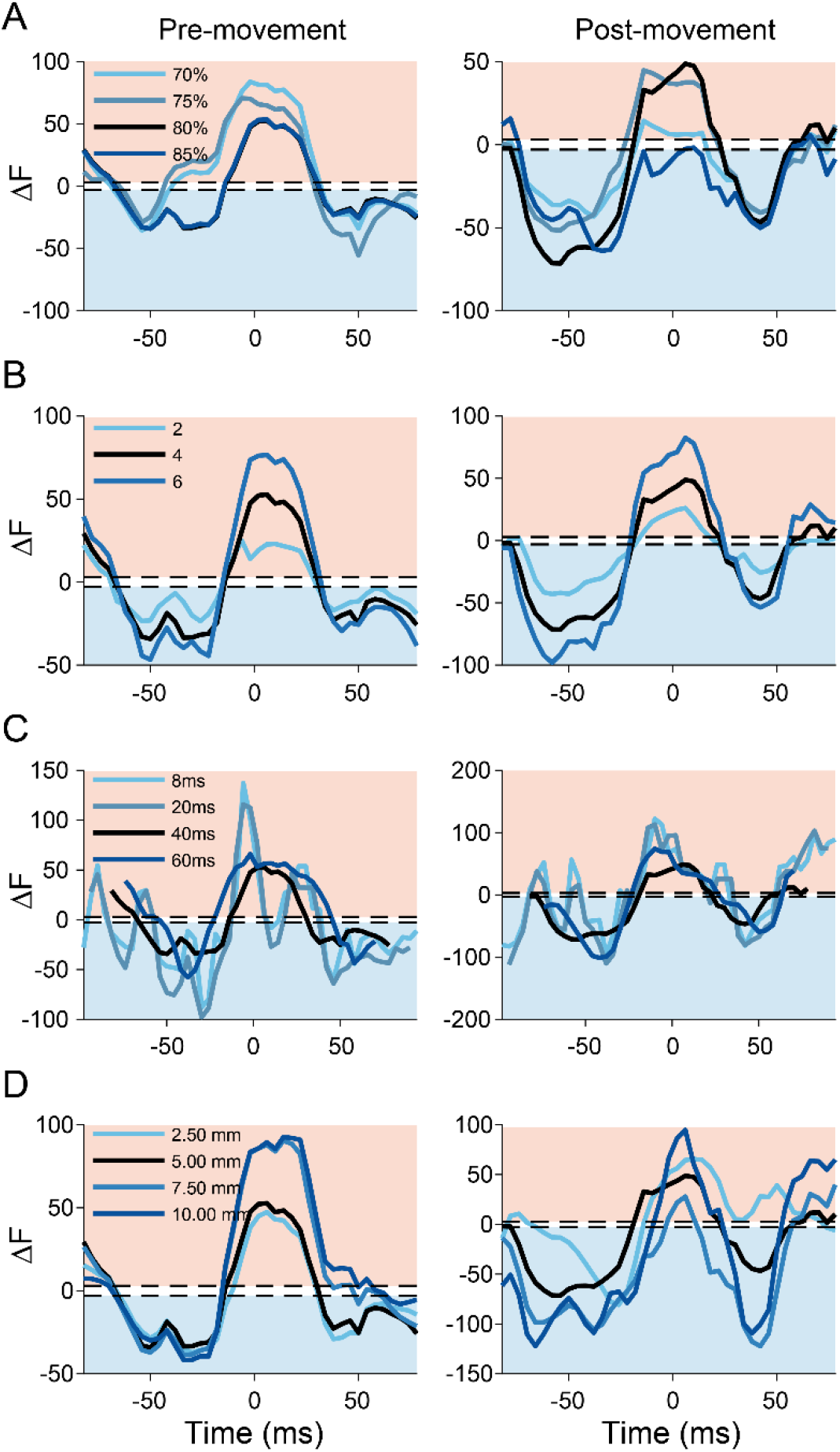
Results with unaligned bursts are robust to analysis parameters. As in Figure S4, for unaligned premovement (left column) and post-movement (right column) bursts.

**Figure S6.**
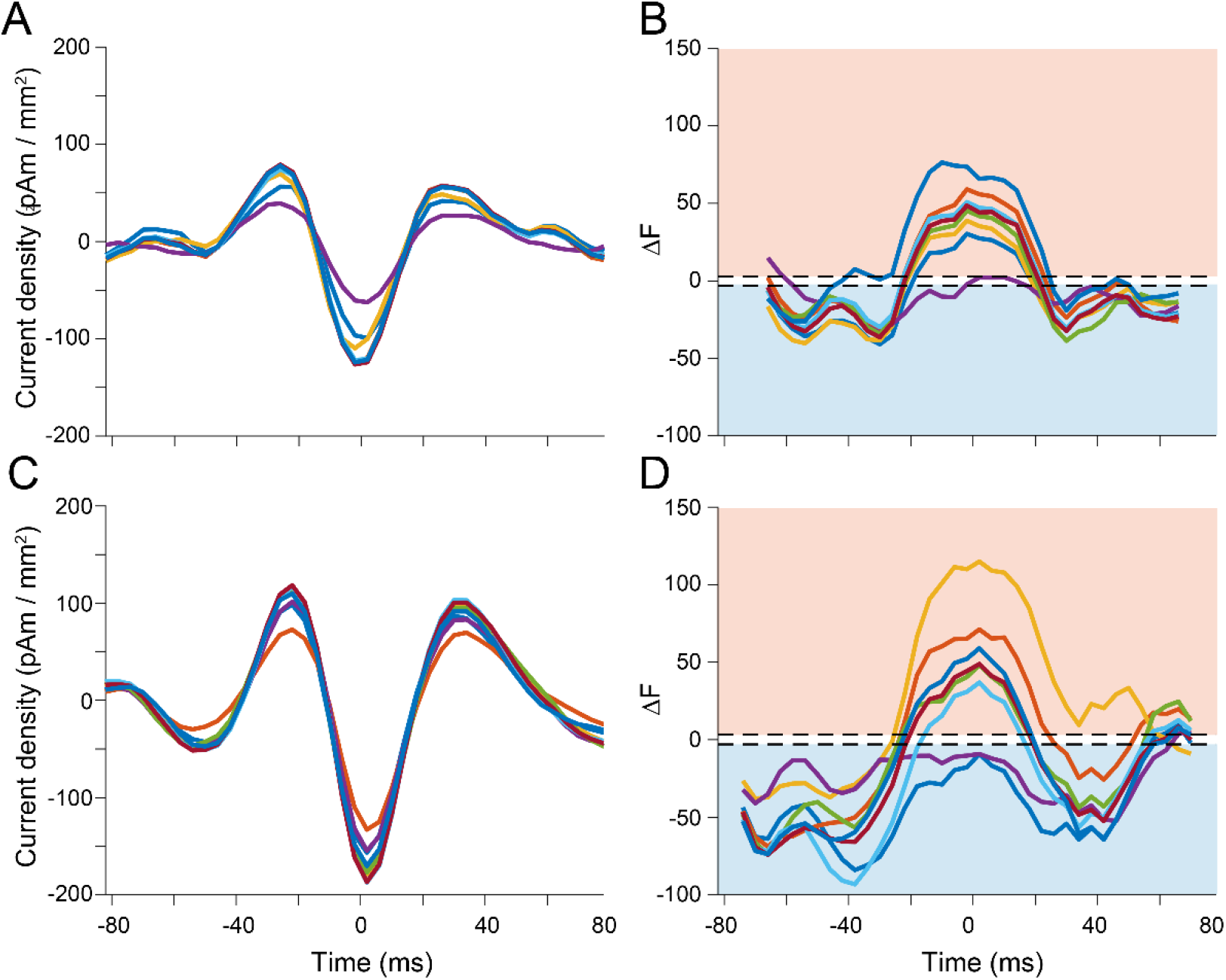
Results are robust to individual subject differences. A) Mean pre-movement beta burst source level current density time courses after excluding each subject (each line shows the mean after excluding a different subject). B) The time course of relatively deep or superficial activity was maintained no matter which subject was left out. This was also true for post-movement beta bursts (C, D).

**Figure S7.**
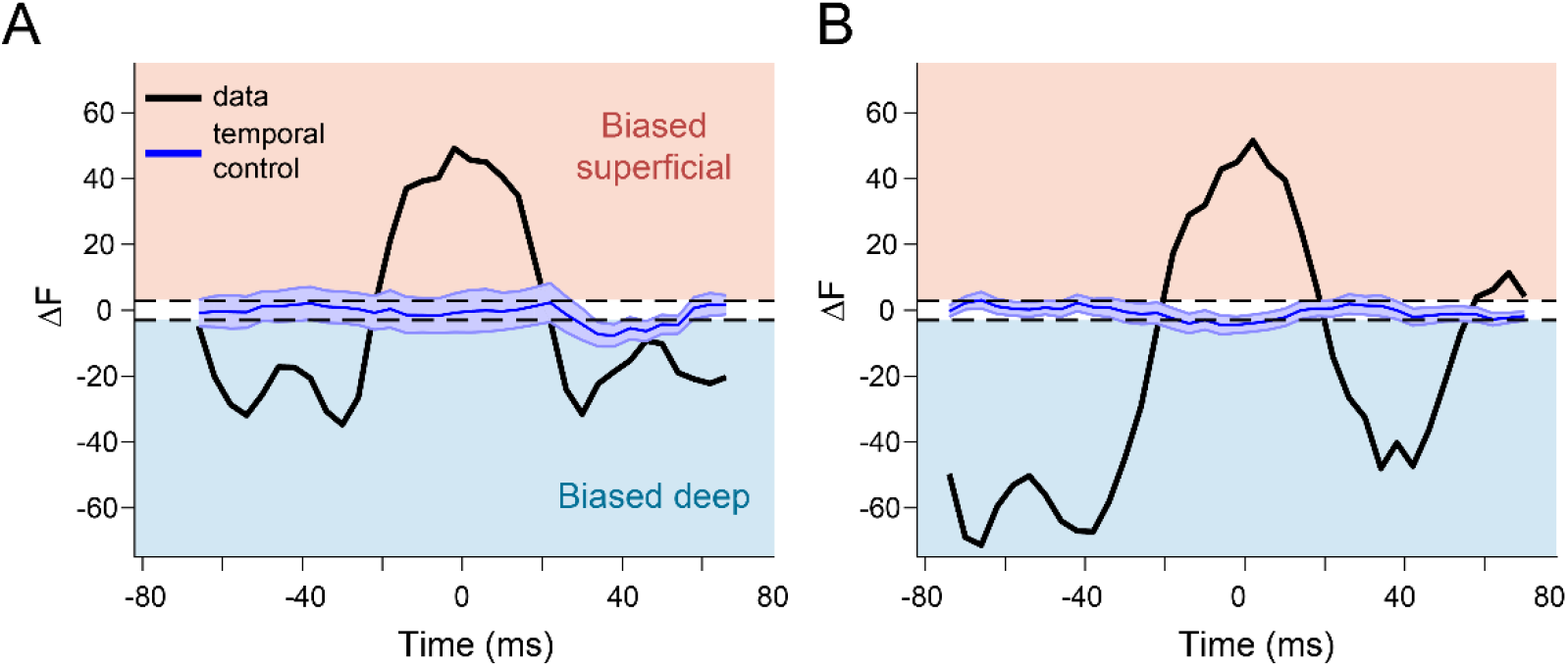
Temporally shuffled surrogate data yields unbiased laminar dominance estimates. Temporally shuffled (blue) surrogate data yields a flat Bayes factor time course that was not biased to either surface for both premovement (A) and post-movement (B) bursts (shaded area shows standard error)

